# Human-induced changes in habitat preference by capybaras (*Hydrochoerus hydrochaeris*) and their potential effect on zoonotic disease transmission

**DOI:** 10.1101/2020.02.05.935445

**Authors:** Thiago C. Dias, Jared A. Stabach, Qiongyu Huang, Marcelo B. Labruna, Peter Leimgruber, Katia M. P. M. B. Ferraz, Beatriz Lopes, Hermes R. Luz, Francisco B. Costa, Hector R. Benatti, Lucas R. Correa, Ana M. Nievas, Patrícia F. Monticelli, Ubiratan Piovezan, Matias P. J. Szabó, Daniel M. Aguiar, José Brites-Neto, Marcio Port-Carvalho, Vlamir J. Rocha

**Affiliations:** Universidade Federal de São Carlos, Rod. Washington Luiz, Km 235, Campus São Carlos, 13565-905, São Carlos, São Paulo, Brasil; Universidade Federal de São Carlos, Rodovia Anhanguera, Km 174, Campus Araras, 13604-900, Araras, São Paulo, Brasil; Conservation Ecology Center, Smithsonian National Zoo & Conservation Biology Institute, Front Royal, Virginia, USA; Faculdade de Medicina Veterinária e Zootecnia, Universidade de São Paulo, Av. Prof. Orlando M. de Paiva, 87, 05508-900, São Paulo, SP, Brasil; Departamento de Ciências Florestais, Escola Superior de Agricultura “Luiz de Queiroz”, Universidade de São Paulo, Av. Pádua Dias, 11, 13418-900, Piracicaba, São Paulo, Brasil; Departamento de Patologia, Programa de Pós Graduação em Biotecnologia do Renorbio, Ponto Focal Maranhão, Universidade Federal do Maranhão, Av. Dos Portugueses, 1966, 65080-805, São Luís, MA, Brasil; Departamento de Patologia, Faculdade de Medicina Veterinária, Universidade Estadual do Maranhão, Av. Lourenço Vieira da Silva, 1000, 65055-310, São Luís, MA, Brasil; Faculdade de Filosofia, Ciências e Letras, Universidade de São Paulo, Av. Bandeirantes, 3900, 14040-900, Ribeirão Preto, São Paulo, Brasil; Embrapa Pantanal, R. 21 de Setembro, 1880, 79320-900, Corumbá, MS, Brasil; Embrapa Tabuleiros Costeiros, Av. Governador Paulo Barreto de Menezes, 3250, 49025-040, Aracaju, SE, Brasil; Laboratório de Ixodologia, Faculdade de Medicina Veterinária, Universidade Federal de Uberlândia, Campus Umuarama, Av. Pará, 1720, 38400-902,Uberlândia, MG, Brasil; Laboratório de Virologia e Rickettsioses, Hospital Veterinário, Faculdade de Medicina Veterinária, Universidade Federal do Mato Grosso, Av. Fernando Corrêa da Costa, 2367, 78060-900, Cuiabá, MT, Brasil; Secretaria Municipal de Saúde, Av. Bandeirantes, 2390, 13478-700, Americana, SP, Brasil; Divisão de Florestas e Parques Estaduais, Instituto Florestal, R. do Horto, 931, São Paulo, SP, Brasil

## Abstract

Human activities are changing landscape structure and function globally, affecting wildlife space use, and ultimately increasing human-wildlife conflicts and zoonotic disease spread. Capybara (*Hydrochoerus hydrochaeris*) is a conflict species that has been implicated in the spread and amplification of the most lethal tick-borne disease in the world, the Brazilian spotted fever (BSF). Even though essential to understand the link between capybaras, ticks and the BSF, many knowledge gaps still exist regarding the effects of human disturbance in capybara space use. Here, we analyzed diurnal and nocturnal habitat selection strategies of capybaras across natural and human-modified landscapes using resource selection functions (RSF). Selection for forested habitats was high across human-modified landscapes, mainly during day- periods. Across natural landscapes, capybaras avoided forests during both day- and night periods. Water was consistently selected across both landscapes, during day- and nighttime. This variable was also the most important in predicting capybara habitat selection across natural landscapes. Capybaras showed slightly higher preferences for areas near grasses/shrubs across natural landscapes, and this variable was the most important in predicting capybara habitat selection across human-modified landscapes. Our results demonstrate human-driven variation in habitat selection strategies by capybaras. This behavioral adjustment across human-modified landscapes may be related to BSF epidemiology.

## Introduction

An increasing number of wild species is being forced to adapt to human-modified landscapes and to live within close proximity to humans [1, 2, 3]. Across these landscapes, human disturbance is linked to shifts in wildlife spatial ecology [4, 5, 6], ultimately affecting zoonosis spread and transmission [7, 8]. In that context, obtain accurate data to address questions on the potential effects of wild species’ habitat use in zoonotic disease transmission is a challenging and crucial goal to wildlife managers and public health institutions.

Capybaras (*Hydrochoerus hydrochaeris*), the largest living rodents on the planet [9], have been rising in human-modified landscapes over the last few decades [10]. These semi-aquatic grazing mammals are usually found in habitats with arrangements of water sources, forest patches and open areas dominated by grasses [11, 12]. Benefited by the great abundance of high-quality food resources from agricultural crops and reduced presence of large predators, capybara populations are recently experiencing a rapid grow [11, 13].

Over some regions, large populations of capybaras are linked to increased crop damage [14], increased vehicle collisions [15], and the spread of Brazilian spotted fever (BSF) - the most lethal spotted fever rickettsioses in the world [16]. Capybaras are responsible for maintaining and carrying large numbers of *Amblyomma sculptum* ticks, the natural reservoir and main vector of the bacterium *Rickettsia rickettsii*, the etiological agent of BSF [16]. Capybaras can also act as amplifying hosts of *R. rickettsii* among *A. sculptum* populations [16, 17]. Even though the role of capybaras in BSF epidemiology have been well-discussed [16, 17, 18, 19], little is known about the potential effects of human-driven variation in capybara habitat selection to BSF spread and transmission.

In this study, we investigated and quantified the variation in diurnal and nocturnal habitat selection strategies by GPS-tracked capybaras across natural and human-modified landscapes. We tested the predictions that: (A) as other mammals (e.g. wildebeest [5]), capybaras must show variation in habitat selection preferences across natural and human-modified landscapes due to different levels of human disturbance in these landscapes; and (B) this variation may be mainly related to temporal avoidance [20, 21] of human activities in human-modified landscapes, with capybaras increasing their selection for forests and water sources during daytime periods.

## Methods

### Ethical statements

Capybara field capture was authorized by the Brazilian Ministry of the Environment (permit SISBIO No. 43259-6), by the São Paulo Forestry Institute (Cotec permit 260108-000.409/2015), and approved by the Institutional Animal Care and Use Committee of the Facult y of Veterinary Medicine of the University of São Paulo (protocol 5948070314).

### Study Area

From 2015 to 2018, we tracked 11 groups of capybaras in Brazil with Lotek Iridium Track M 2D GPS collars (Lotek Wireless, Haymarket, Ontario, CN). Of these, four groups of capybaras were tracked in natural landscapes of Mato Grosso and Mato Grosso do Sul states and seven groups across human-modified landscapes of São Paulo state (Fig 1). To assess the level of human disturbance at our study sites, we incorporated the Human Footprint Index (HFI) from a previous work [22], that ranges from 0 (natural landscapes) to 50 (high-density built landscapes). The spatial resolution of the global dataset is 1-km.

**Fig 1.**
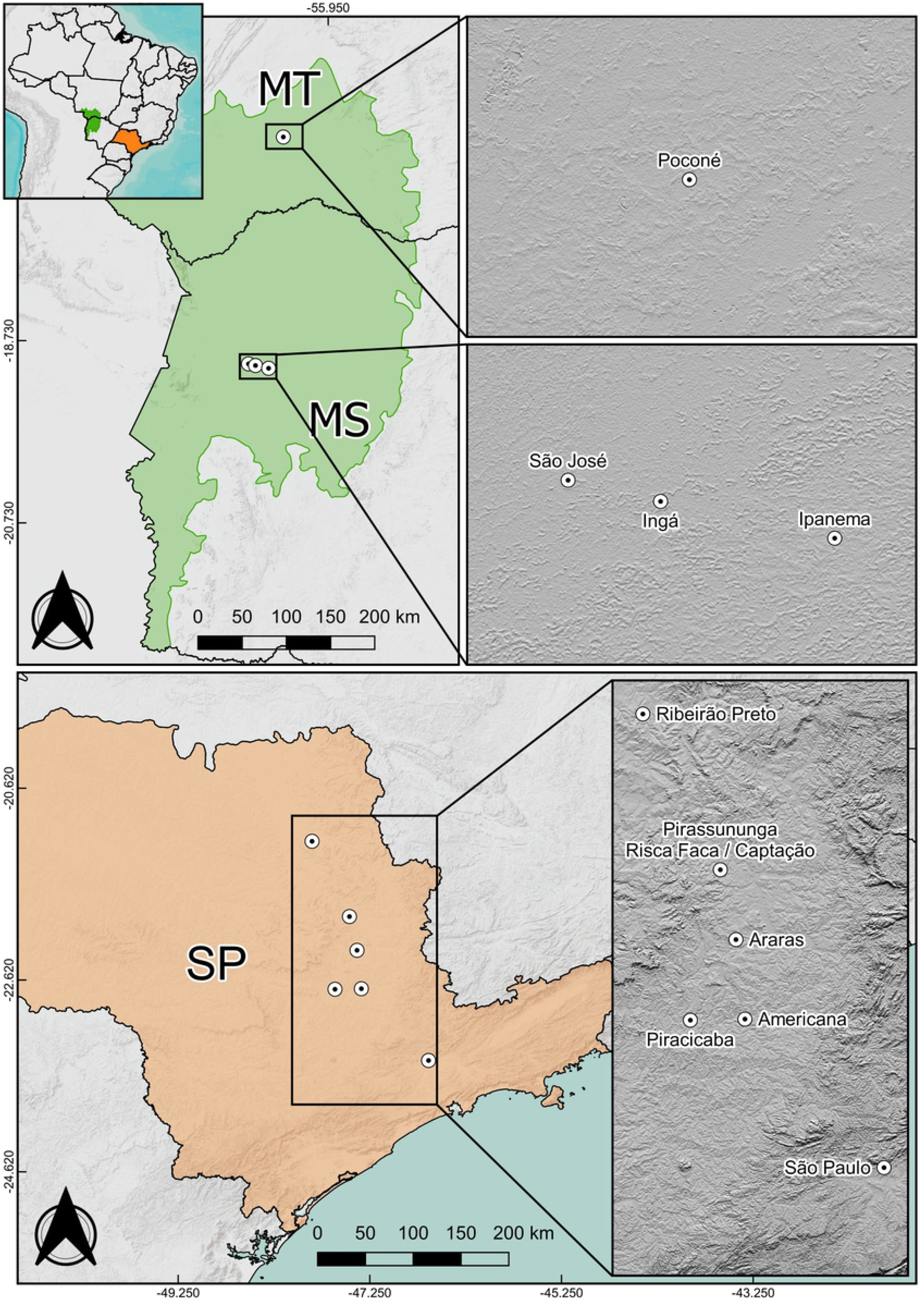
Study areas across natural and human-modified landscapes in Brazil. Study sites in natural landscapes were located at Pantanal biome (green color) in Mato Grosso (MT) and Mato Grosso do Sul (MS) states. Across human-modified landscapes, seven study areas were located in six municipalities in São Paulo state (orange color): Ribeirão Preto, Pirassununga, Araras, Americana, Piracicaba and São Paulo (Geographic Coordinate System: WGS 84 / EPSG 4326).

For all study areas, we summarized the average HFI within the mean dispersal distance of capybaras from their groups (3.4 km) at each centroid [23], calculated using QGIS 2.18.9 [24]. Across natural landscapes, HFI ranged from 2.4 to 6.8 (x̄ = 4.5; n = 4), and in human-modified landscapes the index ranged from 17.4 to 37.7 (x̄ = 29.2; n = 7). In addition, Mato Grosso and Mato Grosso do Sul states have low human population densities when compared to São Paulo state (Mato Grosso = 3 persons km^−2^, Mato Grosso do Sul = 7 persons km^−2^; São Paulo = 166 persons km^−2^) [25].

It is important to emphasize that no case of BSF has been reported in Mato Grosso and Mato Grosso do Sul states, and serological analyses of capybaras from these natural landscapes have shown no evidence of *R. rickettsii* exposure [19]. In contrast, at least three study areas of human-modified landscapes in São Paulo state were classified as BSF-endemic (municipalities of Americana, Araras and Piracicaba), with recent occurrence of human cases and serological evidence of *R. rickettsii* infection in capybaras [19].

Study areas in natural landscapes (São José, Ingá, Ipanema and Poconé) were located in the Pantanal biome. The Pantanal is the largest wetland in the world, characterized by a mosaic of upland vegetation and seasonally flooded areas [26, 27]. This biome consists of large areas of natural vegetation and well-structured/stable ecological communities. The Pantanal support an extraordinary concentration and abundance of wildlife [28], including an impressive assemblage of medium and large carnivores [29, 30]. Within the sampled areas of Pantanal, capybaras had no access to crops or exotic grasses.

Unlike natural landscapes, human-modified landscapes in São Paulo state underwent significant land use and cover changes during the second half of the 19^th^ and early 20^th^ century, transforming natural vegetation (Atlantic rainforest and Cerrado biomes) into a mosaic comprised of small forest fragments surrounded by an agro-pastoral matrix [31]. These forest fragments likely experience large edge effects and reduced biodiversity [32], which affects the abundance of medium and large carnivores across the region. Jaguar (*Panthera onca*), puma (*Puma concolor*), anacondas (*Eunectes* spp.), and caimans (*Caiman* spp.) face threats in the state according to the “São Paulo State Redbook of Fauna Threatened by Extinction” [33].

Across human-modified landscapes, we tracked capybaras in six municipalities: Americana, Araras, Piracicaba, Pirassununga, Ribeirão Preto and São Paulo (Fig 1). With the exception of the municipality of São Paulo, all five areas were located in agricultural landscapes. Sugar cane, corn, cultivated pasturelands, and small forest fragments were the dominant landscape components in the study sites. In Ribeirão Preto, the area used by capybaras was surrounded by a fence that prevented animals from accessing agricultural crops, but they did have access to exotic grasses, as it was also the case in the other human-modified landscapes. In São Paulo municipality, capybaras were monitored in Alberto Löfgren State Park, a protected area within a forest/urban matrix.

### Capybara capture and collaring

In São José, Ingá and Ipanema ranches (natural landscapes), individuals were tranquilized and captured with the aid of a pneumatic rifle (Dan-Inject model JM Standard, Denmark). We used a mixture of ketamine (10 mg/kg) and xylazine (0.2 mg/kg) to anesthetize captured animals [34]. As capybaras use water [11], we targeted animals at a large distance (>20m) from this resource to reduce risk of drowning during tranquilization and capture. Across all other study areas, we captured capybaras through corral-type traps, similar to previously described trap [35].

To better understand movement of capybara populations and minimize the mortality risk of tracked animals, we focused GPS collaring entirely on females. Females show lower agonistic interaction rates when compared to males [36] and therefore, have a decreased chance of mortality. Most female capybara are found in social groups [10, 37] and are thought to be philopatric [38]. We targeted the largest females within each group for GPS collaring because there is a significant correlation between weight and hierarchical position [36]. Hence, we assumed that dominant female movement provided the best representation of group movement.

To avoid incorporating geolocations with large spatial errors [39], we removed GPS positions with a Dilution of Precision (DOP) > 9, following recommendations in Lotek’s GPS collaring manual (Lotek Wireless, Haymarket, Ontario, CN.). The day of capture was removed from analyses to reduce bias in space use related to capture-induced stress [40]. Individuals with < 100 data points were also removed. GPS data were rarified to a 4-hour time interval and categorized into diurnal and nocturnal according to sunrise and sunset time using the ‘*maptools’* package [41] in the R statistical environment [42].

### Habitat data

To generate covariate data for our habitat selection analysis, we performed a supervised land cover classification using Random Forests, an ensemble learning method common for classifying satellite imagery [43]. We used multispectral high-resolution imagery (2-m resolution) acquired by the WorldView-2 satellite (DigitalGlobe, Inc.) and ancillary data derived from each satellite scene for classification (Table A in Appendix S1). We established four habitat classes across natural landscapes (forest, water, grasses/shrubs, bare soil) and five in human-modified landscapes (we added a settlements/roads class). The land cover classification was performed using the ‘*RStoolbox’* package [44] in the R statistical environment.

We digitized 1531 training polygons in QGIS 2.18.9 [24] based on visual interpretation of Worldview-2 satellite scenes. Polygons were divided into calibration (70%; used as input for the land cover classification) and validation (30%; used to evaluate the classification). Overall accuracy ranged from 0.95 to 1 in natural landscapes (*x̄* = 0.97; n = 3) and from 0.84 to 0.99 in human-modified landscapes (*x̄* = 0.94; n = 6). We also applied a post-classification filter to reduce ‘salt-and-pepper’ noise generated by per-pixel classifiers [45]. More details on the land cover classification can be found in Appendix S1.

For each study area, we calculated the Normalized Difference Vegetation Index (NDVI) [46], and created a binary classification of three habitat layers with ecological relevance to capybaras: forest, water and grasses/shrubs. Forest layers included all the types of forested vegetation, primary or secondary, native or not. Water layers included lakes, ponds, and rivers. Grasses/shrubs layers included native and exotic underbrush and shrubby vegetation, including pasturelands, and agricultural crops.

Using binary habitat classifications, we generated distance layers and calculated the shortest distance between each capybara tracking location and habitat classes. For forest distance calculations, we excluded 50-m from the forest edge to assess selection for areas into the forest interior and edges as well. Large double-lane highways found at some of our study sites (varying from 32 to 44 m width: Rodovia Ernesto Paterniani, Rodovia Luis de Queiroz and Rodovia Anhanguera) likely present barriers to capybara’s movement. Because tracked animals did not cross highways during our study, habitats located beyond these highways were not included in our models. Distance to forest interior, distance to water, distance to grasses/shrubs, and NDVI were used as input parameters for resource selection models.

### Resource selection functions

We evaluated habitat selection by comparing the use and availability of habitats through a fine-scale third/fourth-order [47] resource selection function (RSF) analysis [48]. Day and nighttime periods were analyzed separately, due to recognition that capybara habitat use varies throughout the circadian cycle [49]. Habitat availability was determined using a set of random points generated within a predetermined area, as used by other colleagues [5]. We created buffers around each GPS locations with a radius equal to the maximum step length displaced by each animal over a 4-hour period.

To determine the appropriate number of random points per ‘use’ point (GPS location), we performed a sensitivity analysis following details described by previous works [5, 50]. We randomly selected one individual from each study area and fit multiple logistic regression models across several possibilities (1, 2, 3, 5, 10, 20, 30 and 50) of random points. We repeated the process 100 times and calculated the expectation of the coefficient estimates and the 95% simulation envelopes. We determined that a sample of 30 availability points per ‘use’ point provided stable coefficient estimates (Fig A in Appendix S2). The analysis was performed in R [42].

We included habitat variables in our RSF after determining that they were not highly correlated (Pearson’s r > 0.65). To facilitate comparisons across landscapes and cross-time periods, we scaled and centered all data layers ([*x̄* ‒ *x̄*]/*σ*_*x*_). We included quadratic terms for all habitat variables to test for non-linear relationships. Habitat selection was modeled applying a generalized linear mixed-effects logistic regression, following the equation:

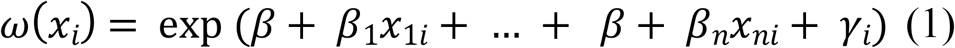

Where *ω*(*x*_*i*_) is the RSF, *β*_*n*_ is the coefficient for the *n*th predictor habitat variable *x*_*n*_, and *γ* is the random intercept for the animal *i*. We incorporated random effects into the model structure to better account for differences between individuals, while also accounting for unbalanced sampling designs [51]. We used nested random effects (“individual” inside “study area” inside “landscape”) to evaluate landscape-level coefficients. A hierarchical approach was used to account for non-independence between individual movements [5]. Habitat selection was modelled using the *‘lme4’* package [52].

### Models

We created four candidate models (forest, water, open areas and full) for each landscape and time-period (Table 1) and used Akaike’s Information Criterion (AIC) to rank them [53]. Models were created to evaluate the importance of different resources on capybara habitat selection: (1) forest - providing shelter from daytime heat and a resting place during the night [26]; (2) water - used by capybaras for thermoregulation, mating and as a refuge from predator attacks [11]; and (3) open areas - used for grazing to meet energy demands [49]. A fourth model, inclusive of all variables, was tested to evaluate if a combination of factors most influenced capybara habitat selection.

**Table 1.**
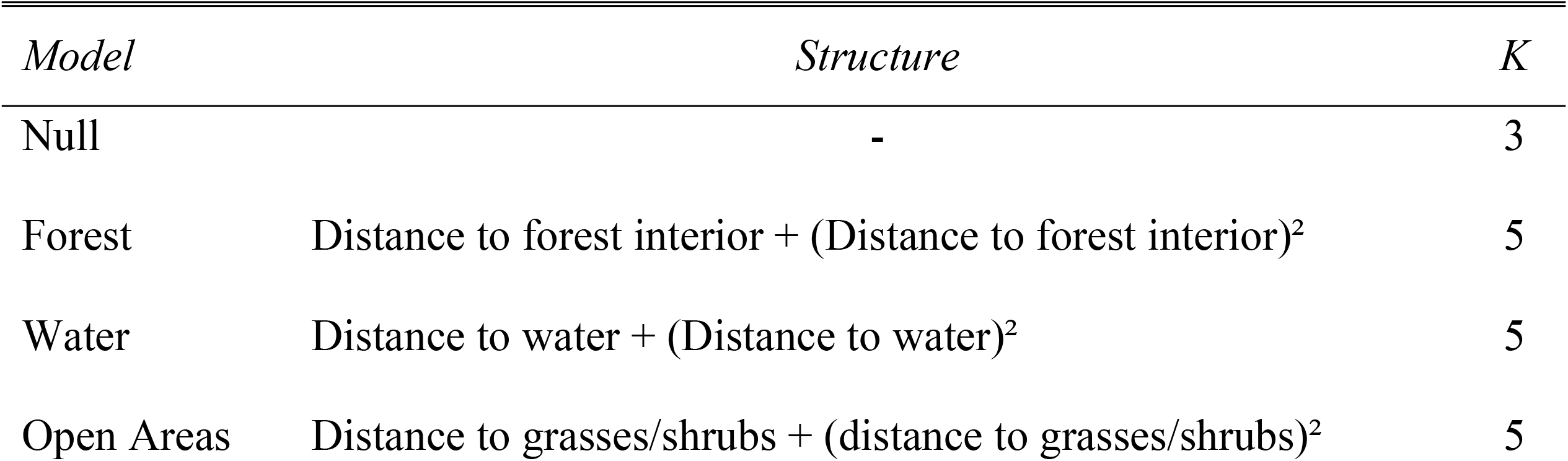
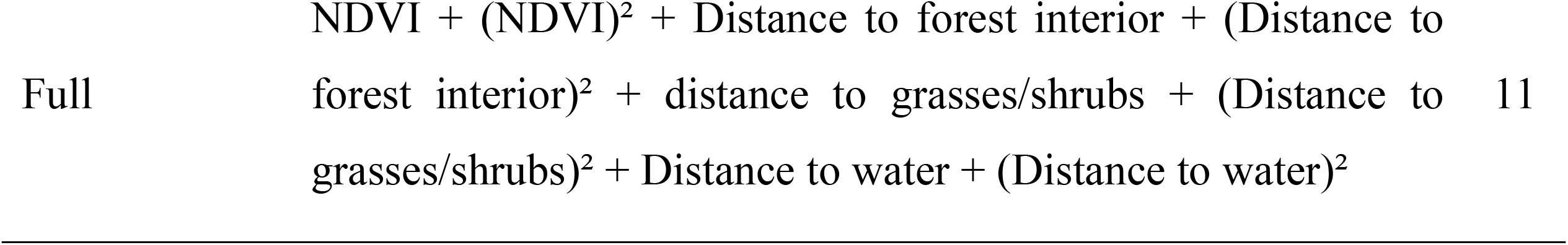
Model structure and number of input variables (K).

We compared all models to a null model using chi-squared tests in R [42]. Coefficients of top-ranked models with confidence intervals that overlap zero were considered statistically insignificant. Top-ranking models were evaluated following the technique in [54], applying Spearman rank correlations between area adjusted frequencies, using presence-only validation predictions and RSF bins (Appendix S3).

## Results

### Capybara capture and collaring

A total of 20 capybaras were captured and fitted with GPS collars. Capybaras were monitored for 33 to 918 days (*x̄* = 273 days), with a similar number of positions collected across study areas (Table S1). Average fix success was high for both landscapes, ranging from 87% to 99% in natural landscapes (*x̄* = 94%;*n* = 4) and from 94% to 99% in human-modified landscapes (*x̄* = 98%;*n* = 16). Maximum distance displaced by individuals in 4-hour time interval ranged from 442-m to 1437-m across natural landscapes (*x̄* = 958.2;*n* = 4) and 268-m to 2703-m in human-modified landscapes (*x̄* = 867.6;*n* = 16).

### Natural landscapes models

The full model was top-ranked across day- and nighttime periods in natural landscapes, indicating that all habitat variables were important in predicting capybara habitat selection (Table 2). Cross-validation highlighted a strong fit to our data (Table A in Appendix S3), with stronger results for daytime periods (*day average r*_*s*_ = 0.83; *night average r*_*s*_ = 0.69). In natural landscapes, distance to water was the most important variable predicting capybara habitat selection (Table 3), with higher coefficient during nighttime periods (day: β = ‒ 1.52 ± 0.03;night: β = ‒ 1.91 ± 0.03; Table 3). NDVI was a weak variable in predicting capybara habitat selection during day periods and was not significant during nighttime periods (day: β = 0.21 ± 0.02;night: β = 0 ± 0.02; Table 3).

**Table 2.**
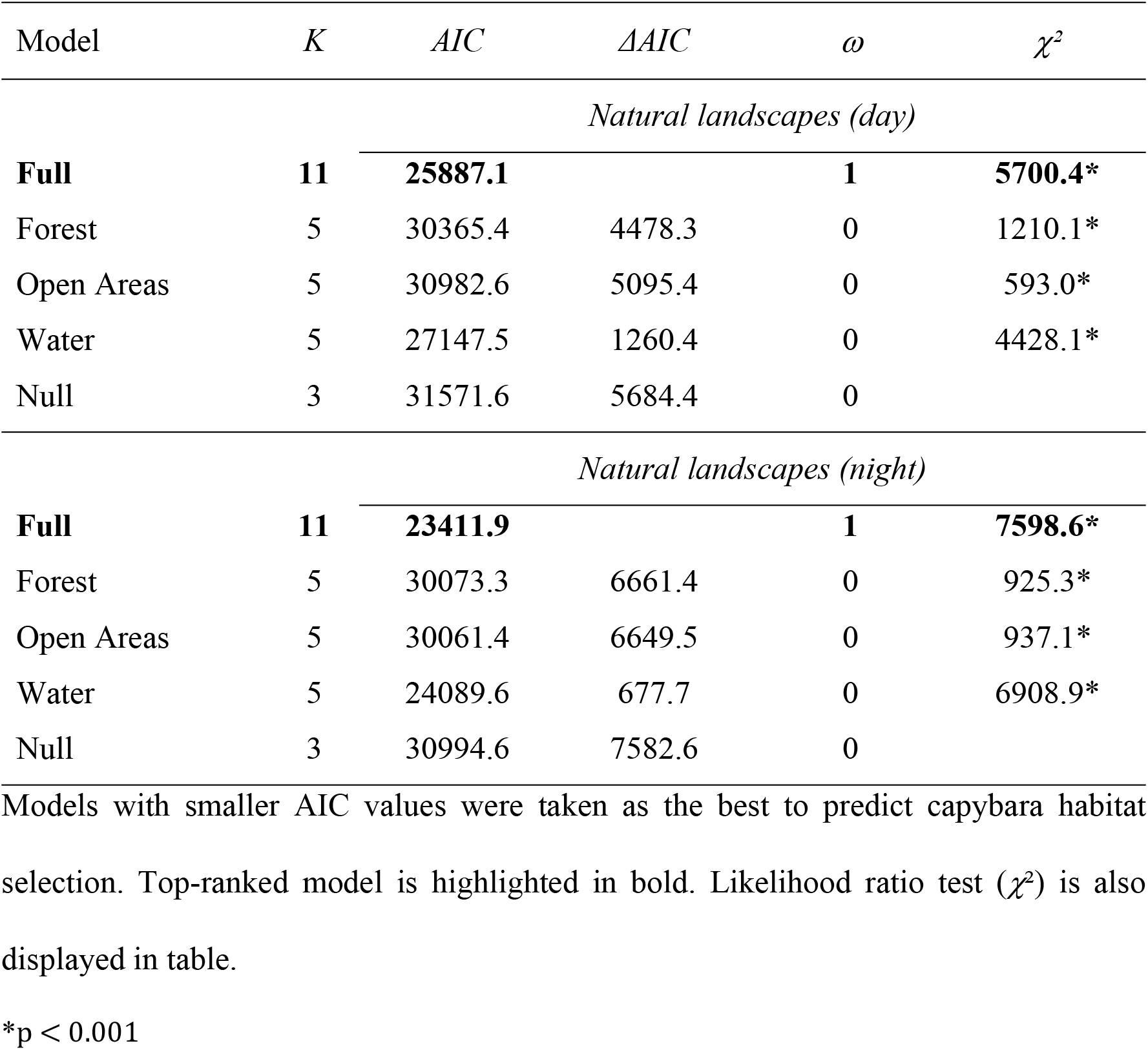
Model selection across natural landscapes for day- and night periods, based on Akaike Information Criterion (AIC).

| Model | $K$ | $AIC$ | $\Delta AIC$ | $\omega$ | $\chi^2$ |
| --- | --- | --- | --- | --- | --- |
| *Natural landscapes (day)* |  |  |  |  |  |
| **Full** | **11** | **25887.1** |  | **1** | **5700.4\*** |
| Forest | 5 | 30365.4 | 4478.3 | 0 | 1210.1* |
| Open Areas | 5 | 30982.6 | 5095.4 | 0 | 593.0* |
| Water | 5 | 27147.5 | 1260.4 | 0 | 4428.1* |
| Null | 3 | 31571.6 | 5684.4 | 0 |  |
| *Natural landscapes (night)* |  |  |  |  |  |
| **Full** | **11** | **23411.9** |  | **1** | **7598.6\*** |
| Forest | 5 | 30073.3 | 6661.4 | 0 | 925.3* |
| Open Areas | 5 | 30061.4 | 6649.5 | 0 | 937.1* |
| Water | 5 | 24089.6 | 677.7 | 0 | 6908.9* |
| Null | 3 | 30994.6 | 7582.6 | 0 |  |

Models with smaller AIC values were taken as the best to predict capybara habitat selection. Top-ranked model is highlighted in bold. Likelihood ratio test ( $\chi^2$ ) is also displayed in table.
\*p < 0.001

**Table 3.** Capybara resource selection function coefficients (β) for both day- and nighttime across natural and human-modified landscapes.

|  | *Natural landscapes* |  | *Human-modified landscapes* |  |
| --- | --- | --- | --- | --- |
|  | *Day* | *Night* | *Day* | *Night* |
| NDVI | **0.21 (0.02)** | 0 (0.02) | **0.32 (0.02)** | 0 (0.02) |
| (NDVI)<sup>2</sup> | -0.01 (0.01) | **-0.03 (0)** | -0.01 (0.01) | **-0.15 (0.01)** |
| Forest Interior | **-0.63 (0.04)** | **-0.32 (0.04)** | **-0.83 (0.04)** | **-0.08 (0.03)** |
| (Forest Interior)<sup>2</sup> | **-0.8 (0.04)** | **-0.72 (0.04)** | **0.21 (0.01)** | **-0.04 (0.01)** |
| Grasses/Shrubs | **0.21 (0.05)** | 0.02 (0.04) | **1.03 (0.03)** | **0.57 (0.03)** |
| (Grasses/Shrubs)<sup>2</sup> | **-0.11 (0.02)** | **-0.02 (0.01)** | **-0.36 (0.02)** | **-0.39 (0.02)** |
| Water | **-1.52 (0.03)** | **-1.91 (0.03)** | **-0.84 (0.02)** | **-0.46 (0.02)** |
| (Water)<sup>2</sup> | **0.32 (0.02)** | **0.66 (0.02)** | **0.16 (0.01)** | -0.01 (0.02) |

Standard errors are displayed within the parentheses; Regression coefficients ( $\beta$ ) with confidence intervals that did not overlapped zero are highlighted in boldface.

Capybaras selected areas further from forest interiors in natural landscapes (Fig 2), with highest probabilities of selection found in areas >250-m from the forest centroid (day- and nighttime periods). Capybaras displayed strong preferences for areas near water. This trend was consistent across day- and nighttime periods (Fig 2), with the probability of selection declining with increasing distance. Preferences for areas near open areas, dominated by grasses/shrubs, were also recorded, with probability of selection decreasing sharply at short distances (Fig 3). Probability of selection by capybaras has increased with increasing NDVI during day- and nighttime periods, although the relatively probability of selection plateaued at a NDVI value of approximately 0.5 during nighttime periods.

**Fig 2.**
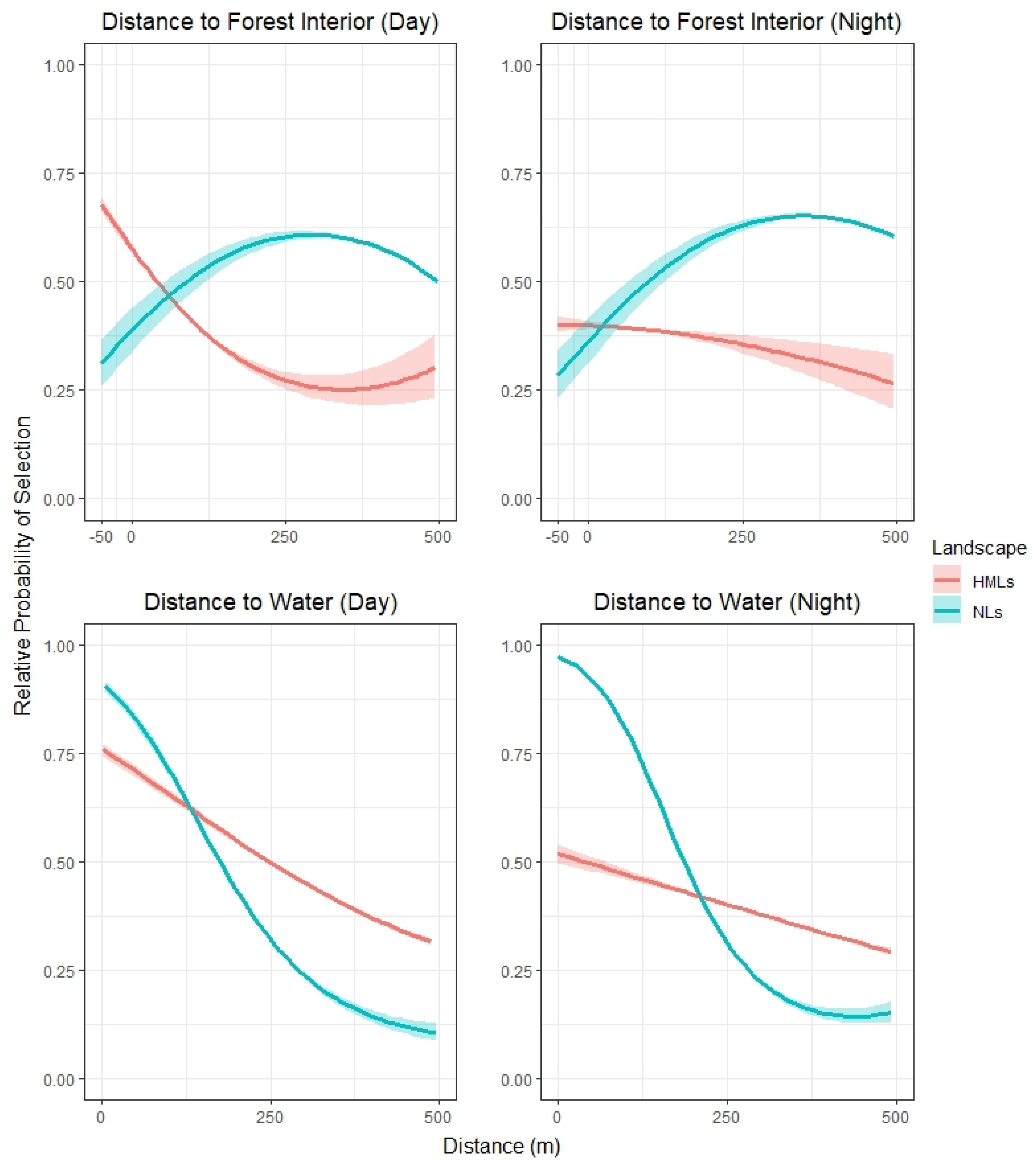
Relative probability of selection of distance to forest interior and distance to water across natural and human-modified landscapes during day- and night periods. The y axis represents the relative probability of selection, ranging from 0 to 1. The x axis represents distance to the habitat. Negative values of forest graphs are related to areas into the forest interior (−50m represents areas 50m inside forest patches).

**Fig 3.**
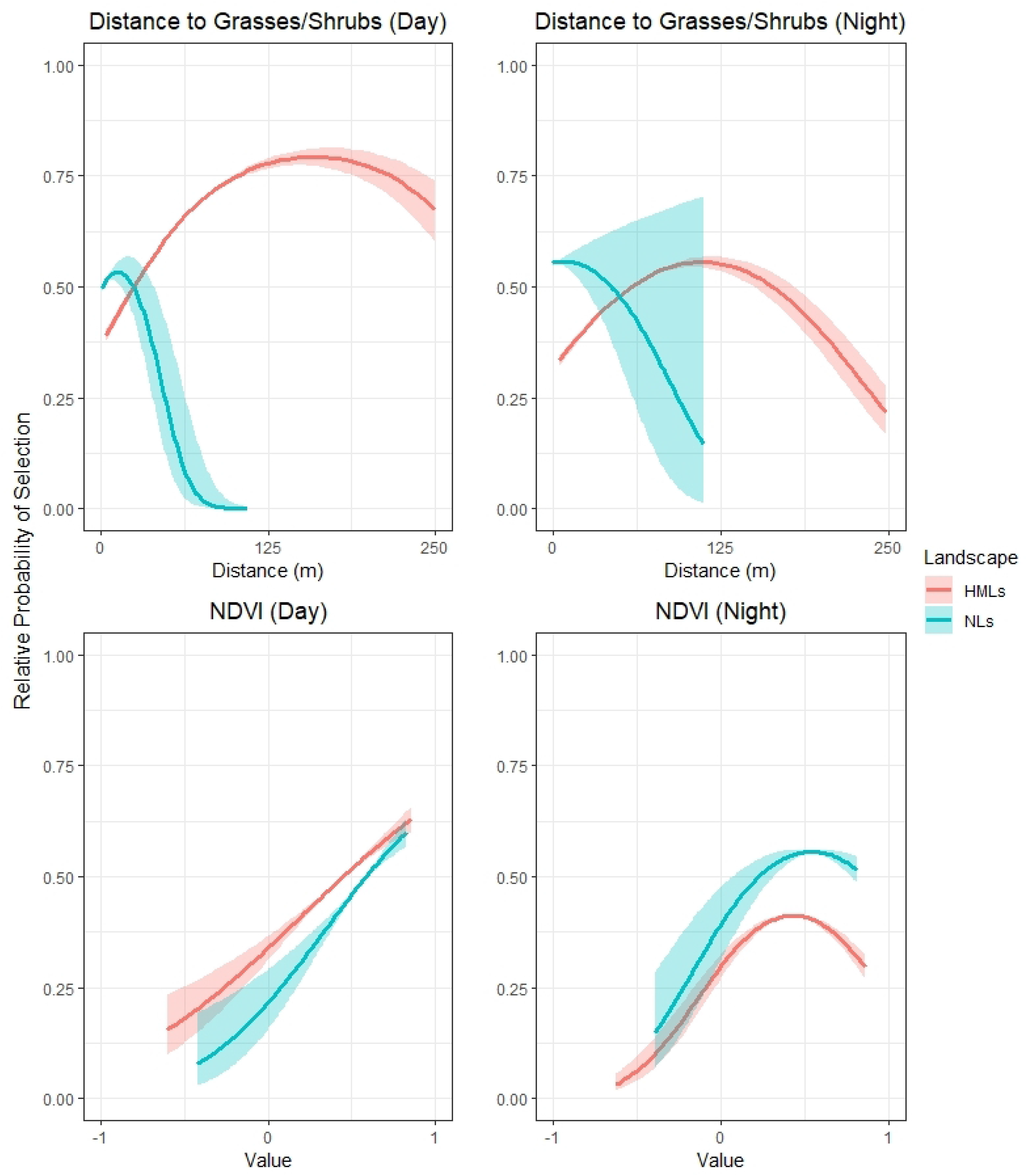
Relative probability of selection for distance to grasses/shrubs and NDVI across natural landscapes and human-modified landscapes during day- and night periods. The y axis represents the relative probability of selection, ranging from 0 to 1. The x axis represents the distance to grasses/shrubs or NDVI values.

### Human-modified landscapes models

Across human-modified landscapes, the full model was also top-ranked for both day- and nighttime periods (Table 4). Models strongly fit the data in these landscapes (*day average r*_*s*_ = 0.89; *night average r*_*s*_ = 0.72), with weaker results found in São Paulo municipality during nighttime, where capybaras were tracked in a non-agricultural state park (Table A in Appendix S3). The most important variable in predicting capybara habitat selection for day- and nighttime periods was distance to grasses/shrubs (day: β = 1.03 ± 0.03;night: β = 0.57 ± 0.03; Table 3). Distance to water (day: β = – 0.84 ± 0.02;night: β = ‒ 0.46 ± 0.02; Table 3) and distance to forest interior (day: β = ‒ 0.83 ± 0.04;night: β = ‒ 0.08 ± 0.03; Table 3) were also significant in predicting capybara habitat selection, with stronger coefficients found for daytime periods. NDVI was a weaker variable in predicting capybara habitat selection during daytime periods, when compared to other habitat variables, and was not significant during nighttime periods (day: β = 0.32 ± 0.02;night: β = 0 ± 0.02; Table 3).

**Table 4.** Model selection across human-modified landscapes for day- and night periods, based on Akaike Information Criterion (AIC).

| Model | *K* | *AIC* | $\Delta AIC$ | $\omega$ | $\chi^2$ |
| --- | --- | --- | --- | --- | --- |
| *Human-modified landscapes (day)* |  |  |  |  |  |
| **Full** | **11** | **40628.2** |  | **1** | **6678.9\*** |
| Forest | 5 | 43675.1 | 3046.9 | 0 | 3620.0* |
| Open Areas | 5 | 43203.9 | 2575.7 | 0 | 4091.2* |
| Water | 5 | 45905.3 | 5277.1 | 0 | 1389.8* |
| Null | 3 | 47291.1 | 6662.9 | 0 |  |
| *Human-modified landscapes (night)* |  |  |  |  |  |
| **Full** | **11** | **44548.5** |  | **1** | **259.5\*** |
| Forest | 5 | 45984.3 | 1435.8 | 0 | 259.5* |
| Open Areas | 5 | 45396.3 | 847.8 | 0 | 847.5* |
| Water | 5 | 45571.3 | 1022.8 | 0 | 672.5* |
| Null | 3 | 46239.8 | 1691.3 | 0 |  |

Models with smaller AIC values were taken as the best to predict capybara habitat selection. Top-ranked model is highlighted in bold. Likelihood ratio test ( $\chi^2$ ) is also displayed in table.
\*p < 0.001

Contrasting to natural landscapes, capybaras across human-modified landscapes were observed with higher preferences for forest interior areas and areas close to forests, with probability of selection declining with increasing distance to forested habitats (Fig 2). Capybaras also showed preferences for areas near water sources, with higher selection during the day (Fig 2). Lower preferences for areas close to grasses/shrubs were found for human-modified landscapes when compared to natural landscapes, with selection increasing at mid distances (125-m) and declining at larger distances (250-m; Fig 3). Similar to natural landscapes, the relative probability of selection increased with increasing NDVI values during daytime periods (maximum coefficients at NDVI values close to 0.7). For nighttime periods, the relative probability of selection peaked at a NDVI value close to 0.5.

## Discussion

This is the first study using GPS tracking, high-resolution imagery and resource selection functions (RSF) to analyze and quantify capybara habitat selection strategies across natural and human-modified landscapes. Capybaras strongly selected forested habitats across human-modified landscapes, which may be a direct response to human activities (e.g. agricultural machinery, people and vehicle traffic), more pronounced during day periods in open areas of our study sites. As wildlife respond to human disturbance following the same principles used by prey encountering predators [55], capybaras may increase their selection for forests during daytime to avoid contact with humans. This behavioral adaptation may be a key point in BSF epidemiology.

Forests are preferred ecological niche of *A. sculptum* ticks [19, 56, 57], the main vectors of the BSF agent (*R. rickettsii*) [16]. In particular, environmental tick burdens were found to be much higher across human-modified landscapes than across natural landscapes studied here [19]. Therefore, capybaras may be highly efficient hosts across human-modified landscapes, increasing their capacity in maintaining and carrying large numbers of *A. sculptum* [16, 19], due to shared preferences for forested habitats.

As efficient vertebrate hosts for *A. sculptum* across human-modified landscapes, capybaras are linked to BSF spread due to increased chance of infection by *R. rickettsii* and translocation of infected ticks. Capybaras are also linked to the amplification of rickettsial infection among *A. sculptum* populations, creating new cohorts of infected ticks during bacteremia periods (days or weeks), when they maintain *R. rickettsii* in their bloodstream [16]. In addition, *A. sculptum* populations are not able to sustain *R. rickettsii* for successive generations without the creation of new infected cohorts via horizontal transmission through vertebrate hosts [58, 59]. Therefore, we highlight the role of capybaras selecting disturbed forest in human-modified landscapes as an important factor in BSF spread.

Preferences for areas nearby water sources across natural and human-modified landscapes were not surprising. Capybaras are semi-aquatic mammals and their dependence on water sources has already been well-documented, with some authors reporting these rodents hardly moving more than 500-m from water [60, 61]. However, our models highlighted that capybaras were less dependent on water sources in human-modified landscapes, which may be related to human-driven variation in one or more behaviors linked to water use: reproduction, thermoregulation, or predator avoidance [11].

Quality and quantity of food resources from highly nutritious agricultural and pasture fields seems to have a strong influence in habitat selection by, since grasses/shrubs was the strongest variable in our human-modified landscapes’ models. Because we wanted to compare selection for similar habitats across natural and human-modified landscapes, we did not separate crops and pastures into individual habitat classes. However, in the future, more detailed habitat selection studies for capybaras might want to consider fine-scale spatiotemporal dynamics of agriculture and pasture fields in human-modified landscapes. Understanding selection for these resources, mainly sugar cane, which is linked to the BSF spread [62], may be essential to develop conflict mitigation strategies for the species.

Lastly, improving NDVI temporal resolution could potentially increase the link between this vegetation index and capybaras, since this variable was weak in predicting capybara habitat selection. Higher temporal resolution of NDVI may allow for further investigations on the interaction between vegetation quality and capybara habitat use.

Increasingly, wildlife is forced to adapt to human-modified landscapes and live within proximity to humans [3]. Capybaras appear to be well adapted to anthropic environments, with increased abundance and broadened distribution in Brazil [11]. This is likely due to high availability of nutritious resources from agricultural crops and cultivated exotic grasses, and to lower predation risk in human-modified landscapes [13]. The proximity between wildlife and humans has been shown to lead to increase human wildlife conflicts, including zoonotic disease transfer [8]. Across human-modified landscapes, large groups of capybaras have been linked to increased crop damage [14] and vehicle collisions [15], as well as public health issues related to BSF spread [16].

Our results showed clear distinctions between habitat selection of capybaras in natural and human-modified landscapes, providing a background for further investigation into the potential indirect effects of human disturbance in capybara space use. The development of knowledge regarding these effects may assist future management actions aimed at reducing conflicts linked to the species, including those related to Brazilian spotted fever (BSF) spread.

## Conclusions

Through the use of GPS tracking and resource selection functions it was possible to demonstrate variation in habitat selection strategies of capybaras across natural and human-modified landscapes. Forested habitats were more used through human-modified landscapes than across natural landscapes. In addition, capybaras consistently selected areas near water in both landscapes, but this resource was more important in predicting capybara habitat selection in natural landscapes. In contrast, grasses/shrubs (which includes crops and pasture fields) was a stronger predictor of capybara habitat selection across human-modified landscapes. Our results show the influence of anthropic disturbance in capybara space use patterns. The understanding of capybara habitat use in natural and human-modified landscapes may support human-wildlife conflict management and Brazilian spotted fever spread control.

## Acknowledgments

We are grateful to SESC Pantanal, UFMT, Embrapa Pantanal and Alegria and São José ranches (Corumbá) for logistical support during field work in the Brazilian Pantanal and to the “Departamento de Água e Esgoto de Americana (DAE)” for allowing us to work at the “Estação de Tratamento de Esgoto (ETE) de Carioba” in Americana municipality. Forestry Institute and Coordination of Urban Parks of the São Paulo State Secretariat for the Environment offered logistical support in captures and monitoring (SMA Process 000.409/2015). We are also very grateful to GIS Lab at Smithsonian Institution, VA, USA, for the technical training in GIS and spatial analysis.

## Supporting information captions

**S1 Appendix. Land cover classification of capybara habitats.** Methods on how habitats of studied capybaras were classified using high-resolution satellite imagery and random forest algorithm.

**Appendix S2. Sensitivity Analysis performed for study areas across natural and human-modified landscapes.** We performed sensitivity analysis to set the number of random points per ‘use’ point to our habitat selection models.

**Appendix S3. Top-ranked models’ evaluation.** We used presence-only data to evaluate our top-ranked models’ performance through Spearman rank correlations between area-adjusted frequencies and resource selection functions spatial bins.

**Table S1.** Summary table for GPS-tracked capybaras across natural and human-modified landscapes.

